# Benchmarking workflows to assess performance and suitability of germline variant calling pipelines in clinical diagnostic assays

**DOI:** 10.1101/643163

**Authors:** Vandhana Krishnan, Sowmi Utiramerur, Zena Ng, Somalee Datta, Michael P. Snyder, Euan A. Ashley

## Abstract

Benchmarking the performance of complex analytical pipelines is an essential part of developing Laboratory Developed Assays (LDT). Reference samples and benchmark calls published by Genome in a Bottle (GIAB) Consortium have enabled the evaluation of analytical methods. However, the performance of such methods is not uniform across the different regions of the genome/exome and different variant types and lengths. Here we present a scalable and reproducible, cloud-based benchmarking workflow that can be used by clinical laboratories to rapidly access and validate the performance of LDT assays, across their regions of interest and reportable range, using a broad set of benchmarking samples.

## Background

Next Generation Sequencing (NGS) and analytical methods developed to detect various forms disease-causing polymorphisms are now routinely being used by clinical laboratories to determine the molecular etiology of complex diseases/disorders and in many cases to make critical treatment course decisions. In the past two decades, many polymorphisms in the human genome have been identified and validated that serve as predictive, diagnostic, and prognostic markers for complex inherited diseases. These genomic disease markers can be of different forms such as Single Nucleotide Variants (SNVs), small INsertions and DELetions (INDELs), large deletions and duplications (del/dups), and Copy Number Variations (CNVs) and can vary in size from a single base change to several Mega Bases (MB) in length and even whole chromosomal polysomy. Clinically relevant polymorphisms occur in different regions of the genome, including exonic, splice-sites, and deep-intronic regions. These polymorphisms also happen in various forms, including single base changes within high entropic regions, copy number changes to homopolymer repeats and copy number changes to Short Tandem Repeat (STR) regions. NGS platforms used to detect these polymorphisms; owing to their different sequencing chemistry and signal processing methods; have very different error modes and hence very different analytical performance across the different regions of the genome. Consequently, analytical methods specific to various NGS platforms such as Illumina, Ion Torrent, Pacific Biosciences, and Oxford Nanopore have been developed to both account for and correct the errors particular to these sequencing platforms. This has resulted in a dizzying array of combinations of sequencing platforms and analytical methods available to a clinical diagnostic laboratory to develop their LDT assay.

Benchmarking methods and pipelines are essential to accurately assess the performance of sequencing platforms and analytical methods before they are incorporated into clinical diagnostic assays. Genome In A Bottle (GIAB) consortium hosted by NIST has characterized the pilot genome (NA12878/HG001) (1) and six samples from the Personal Genome Project (PGP) (2). These benchmark calls for SNVs and small INDELs (1-20bp) from reference samples can be used for optimization, performance estimation, and analytical validation of LDT assays using complex analytical pipelines with multiple methods to detect polymorphisms in the genome. Global Alliance for Genomics and Health (GA4GH) benchmarking team have developed standardized tools (3) to evaluate the performance metrics of germline variant callers used primarily in research applications.

Clinical Laboratory Improvement Amendments (CLIA) requires all laboratories using LDT to establish the test’s performance specifications such as analytical sensitivity, specificity, reportable range, and reference range (4). College of American Pathologist (CAP) laboratory standards for NGS based clinical diagnostic (5) not only require the laboratories to assess and document the performance characteristics of all variants within the entire reportable range of LDTs but also obtain the performance characteristics for every type and size of variants that are reported by the assay. Laboratories are also required to assess the performance characteristics for clinically relevant variants such as Δ*F*508 and IVS8-5T (6) mutations in a CFTR assay. CAP guidelines also require laboratories to periodically (determined by the laboratory) assess and document the analytical performance characteristics to ensure that the LDT is continuing to perform as expected over time.

Benchmarking workflows/pipelines that are highly scalable, reproducible and capable of reporting the performance characteristics using a large number of reference and clinical samples within multiple highly stratified regions of interest are essential for clinical laboratories to optimize and routinely assess the performance of their LDT assays.

## Results

Our goal was to develop a benchmarking workflow that any clinical laboratory could use to quickly evaluate and compare the performance characteristics of all suitable secondary analysis pipelines. Benchmarking workflow should further help optimize the analytical pipeline based on well-defined performance metrics and finally produce a thorough analytical validation report to justify the use of the analytical pipeline in their diagnostic assay to regulatory authorities such as CLIA and CAP.

To test the abilities of our benchmarking workflow, we used it to compare two analytical pipelines commonly used for germline variant calling 1. Pipeline based on Broad Institute’s best practices guidelines using GATK HaplotypeCaller v3.7 and 2. SpeedSeq pipeline (7) based on FreeBayes v0.9.10 (8) as the primary variant calling engine. GATK HaplotypeCaller based pipeline was chosen over the FreeBayes pipeline as it out-performed in the detection of small-INDELs (1 – 20 base pairs).

The performance characteristics of the analytical pipeline using GATK v3.7 was further optimized using benchmarking metrics generated using the five GIAB reference samples and four GeT-RM samples (see Methods) with known pathogenic variants. Also, it is critical for the clinical laboratories developing NGS based LDT assays to accurately determine the reportable range to avoid misdiagnosis leading to wrong treatment decisions. To this effect, we evaluated the performance metrics using the benchmarking workflow in three distinct genomic regions of interest (see Methods for details).

Although we have the benchmarking results for the region, including coding exons in all the RefSeq genes, we have omitted those findings in this section and focus on the clinically relevant regions.

Tables 1 and 2 show the benchmarking metrics for SNPs in all five GIAB samples within the clinically relevant genes and whole exome regions, respectively. The precision, recall, and NPA metrics for SNPs are uniform across all the reference samples, and there is no sample bias in the results for some of the better-characterized samples such as NA24385 and NA12878. Performance metrics for SNPs within the clinically relevant gene region is significantly better than those within the whole exome region. Recall metrics, in particular, are a percentage point better in the clinically pertinent gene region, across all reference samples. This is attributable to the fact that many genes have isoforms, resulting in higher alignment errors, and some genes have either very high or very low GC content, resulting in higher than average sequencing errors within these regions of the genome. The finding is of great clinical significance, since the reportable region of most inherited disease/disorder, LDT assay is limited to the clinically relevant genes and thereby the overall performance characteristics of the assay is better than that estimated over either the whole genome or whole exome regions.

**Table 1:**
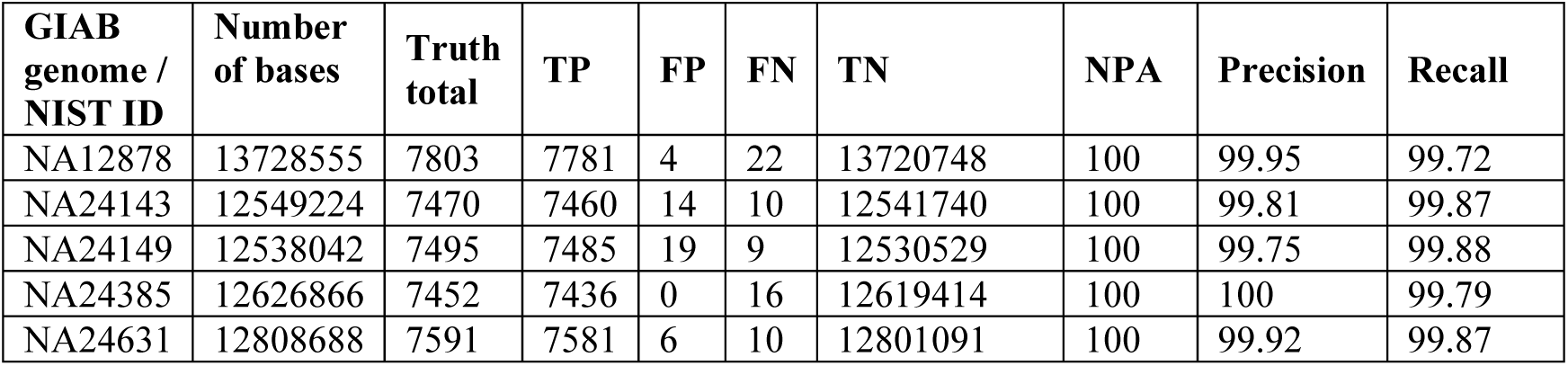
Benchmarking metrics for SNPs within coding exons of clinically relevant ∼7000 genes (as specified in Methods).

| GIAB genome / NIST ID | Number of bases | Truth total | TP | FP | FN | TN | NPA | Precision | Recall |
| --- | --- | --- | --- | --- | --- | --- | --- | --- | --- |
| NA12878 | 13728555 | 7803 | 7781 | 4 | 22 | 13720748 | 100 | 99.95 | 99.72 |
| NA24143 | 12549224 | 7470 | 7460 | 14 | 10 | 12541740 | 100 | 99.81 | 99.87 |
| NA24149 | 12538042 | 7495 | 7485 | 19 | 9 | 12530529 | 100 | 99.75 | 99.88 |
| NA24385 | 12626866 | 7452 | 7436 | 0 | 16 | 12619414 | 100 | 100 | 99.79 |
| NA24631 | 12808688 | 7591 | 7581 | 6 | 10 | 12801091 | 100 | 99.92 | 99.87 |

**Table 2:** Benchmarking metrics for SNPs in whole exome regions, including non-coding exons, splice sites (+/-20 bp) and clinically relevant deep intronic regions.

| GIAB genome / NIST ID | Number of bases | Truth total | TP | FP | FN | TN | NPA | Precision | Recall |
| --- | --- | --- | --- | --- | --- | --- | --- | --- | --- |
| NA12878 | 71152019 | 57822 | 57024 | 491 | 776 | 71093728 | 100 | 99.15 | 98.66 |
| NA24143 | 65657646 | 55975 | 55340 | 669 | 611 | 65601026 | 100 | 98.81 | 98.91 |
| NA24149 | 65597266 | 55518 | 54827 | 669 | 669 | 65541101 | 100 | 98.79 | 98.79 |
| NA24385 | 65948744 | 56068 | 55329 | 389 | 705 | 65892321 | 100 | 99.30 | 98.74 |
| NA24631 | 66988987 | 56948 | 56303 | 394 | 643 | 66931647 | 100 | 99.31 | 98.87 |

Tables 3 and 4 provide the indel benchmarking metrics for sample NA24385 in the clinically relevant and whole exome regions, respectively. As expected, the benchmarking workflow reveals that the performance metrics for INDELs are lower than those for SNPs. However, the stratification by INDEL size, helped us determine the reference range for INDELs (1-20 base-pairs). The recall metric for INDELs larger than 20 base-pairs is significantly lower than the recall for INDELs 1 – 20 base-pairs. As in the case of SNPs, performance metrics for INDEL detection within the clinically relevant genes of interest is better than the whole exome region.

**Table 3:** Benchmarking metrics for indels of different size ranges in NA24385 (truth set NIST v3.3.2, total bases = 12,626,866) for the regions within ∼7000 clinically relevant genes (as specified in Methods).

| Size of indels in NA24385 | Truth total | TP | FP | FN | TN | NPA | Precision | Recall |
| --- | --- | --- | --- | --- | --- | --- | --- | --- |
| 1-10 | 145 | 136 | 12 | 9 | 12626709 | 100 | 91.89 | 93.79 |
| 11-20 | 9 | 9 | 0 | 0 | 12626857 | 100 | 100 | 100 |
| 21-50 | 3 | 3 | 0 | 0 | 12626863 | 100 | 100 | 100 |
| All Indels | 157 | 148 | 12 | 9 | 12626697 | 100 | 92.50 | 94.27 |

**Table 4:** Benchmarking metrics on the number of indels of different size ranges in NA24385 (truth set NIST v3.3, total bases = 65,948,744) for the whole exome regions including non-coding exons, splice sites (+/-20 bp) and clinically relevant deep intronic regions.

| Size of indels in NA24385 | Truth total | TP | FP | FN | TN | NPA | Precision | Recall |
| --- | --- | --- | --- | --- | --- | --- | --- | --- |
| 1-10 | 5169 | 4727 | 872 | 442 | 65942703 | 100 | 84.43 | 91.45 |
| 11-20 | 203 | 188 | 10 | 15 | 65948531 | 100 | 94.95 | 92.61 |
| 21-50 | 67 | 56 | 3 | 11 | 65948674 | 100 | 94.92 | 83.58 |
| All Indels | 5362 | 4920 | 885 | 468 | 65942471 | 100 | 84.75 | 91.27 |

The benchmarking results of the other GIAB reference samples in the clinically relevant and whole exome regions can be obtained in the Supplementary Materials Table S1-S4 and Table S5-S8, respectively. The histogram for the indel size distribution in the NA24385 reference sample for the whole exome region is in Supplementary Material as Fig S1. The histograms of indel size distributions for GIAB samples in both the whole exome and clinically relevant regions are available in our github repository - vandhana/stanford-benchmarking-workflows.

Finally, our benchmarking workflow was able to confirm that our variant calling pipeline can detect all the clinical variants in GeT-RM samples listed in Table 5.

**Table 5:** Validation of the presence of the truth variants in the GeT-RM samples (as specified in Methods) using our variant calling pipeline.

| GeT-RM Sample ID | Chromosome:Position | Truth Variant | Truth Variant Detected |
| --- | --- | --- | --- |
| NA04408 | 15:91310152 | TATC -> T | Yes |
|  | 15:91310156 | T -> TA | Yes |
|  | 15:91310158 | A -> ATTC | Yes |
| NA14090 | 17:41276044 | ACT -> A | Yes |
| NA14170 | 13:32914437 | GT -> G | Yes |
| NA16658 | 10:43609103 | G -> T | Yes |

To get all the metrics produced by hap.py and other output files including plots from our benchmarking workflow for each reference sample, please refer to the Supplementary Data files.

## Discussion

GIAB consortium has helped developed standards for genomic data to evaluate the performance of NGS sequencing platforms and analytical methods used for alignment and variant calling. The precisionFDA platform has enabled the genomics community to develop and deploy benchmarking tools that can evaluate the performance of analytical methods against the gold standard datasets. These benchmarking tools, along with accuracy challenges, has led to the development of highly accurate variant calling methods. However, the requirements of a clinical diagnostic laboratory go beyond the simple evaluation of performance characteristics of an analytical pipeline against one or more reference samples. Our purpose was to build a benchmarking workflow to meet the assay optimization and validation needs of a clinical laboratory. The primary benefit of our benchmarking workflow is that it allows for the assay performance to be evaluated using a broad set of both reference samples with a large number of gold-standard variant calls and clinical samples with a small number of clinical variants, that are specific to the diagnostic assay being evaluated. The benchmarking workflows enable the clinical laboratories to establish the reporting range of the diagnostic assay by estimating the performance within multiple regions of interest.

Unlike web-based benchmarking apps, such as those provided by the precision FDA platform or GA4GH, our benchmarking framework can be seamlessly integrated with any variant calling pipeline in the user’s software environment. Thus, our benchmarking workflows enable ease of use and avoid the transfer of sensitive data to different locations, which could be non-Protected Health Information (PHI) compliant.

Our benchmarking modules if integrated with deployment tools, such as Jenkins (9) and CircleCI (10), that work on the principle of continuous integration and continuous delivery/deployment (CI/CD), it provides a foolproof way of examining consistency in results. In this era where workflows generating reproducible results are gaining attention, easy incorporation of workflows with CI/CD tools is a nice feature to have.

The benchmarking workflow is distributed using human-readable YAML (11) format, and it might limit direct porting to existing WDL based workflows such as those published by the Broad Institute (12, 13). Similarly, conversion of the benchmarking YAML files to Common Workflow Language (CWL) format is required to run workflows published by GA4GH (14-16). However, since we have used docker images for the software tools used within the benchmarking framework, portability to other runtime environments should not take a significant effort for a bioinformatician.

## Conclusions

Benchmarking variants is a critical part of implementing variant calling pipelines for research or clinical purposes. Here, we have successfully implemented benchmarking workflows that generate metrics such as specificity, precision, sensitivity for germline SNPs, and indels in whole exome sequencing data. Also, indel size distributions even in the form of histograms are provided. Combining these benchmarking results with validation using known variants of clinical significance in publicly available cell lines, we were able to establish our variant calling pipelines in a clinical setting. Our benchmarking workflow can serve as a plug-in to any existing variant calling pipeline to work as an integrated unit or be used as a separate module as well.

## Methods

### Benchmarking workflow

The benchmarking workflow, as illustrated in Figure 1, is a sequence of steps required to perform a rapid and comprehensive analytical validation of a clinical diagnostic assay based on germline variants. The benchmarking workflow can be easily integrated with any secondary-analysis pipeline used in a diagnostic assay to call germline variants, and our workflow accepts germline variants (SNVs and small INDELs) in Variant Call Format VCF v4.1(17) or higher. The workflow takes one or more stratification files specifying the regions of interest in BED (18) format and generates a comprehensive analytical validation report detailing the performance characteristics of the assay within each of the specified regions of interest. Benchmark variant calls that are considered as ground truths for each of the reference sample used to evaluate the analytical performance can be also be specified in VCF format.

**Figure 1.**
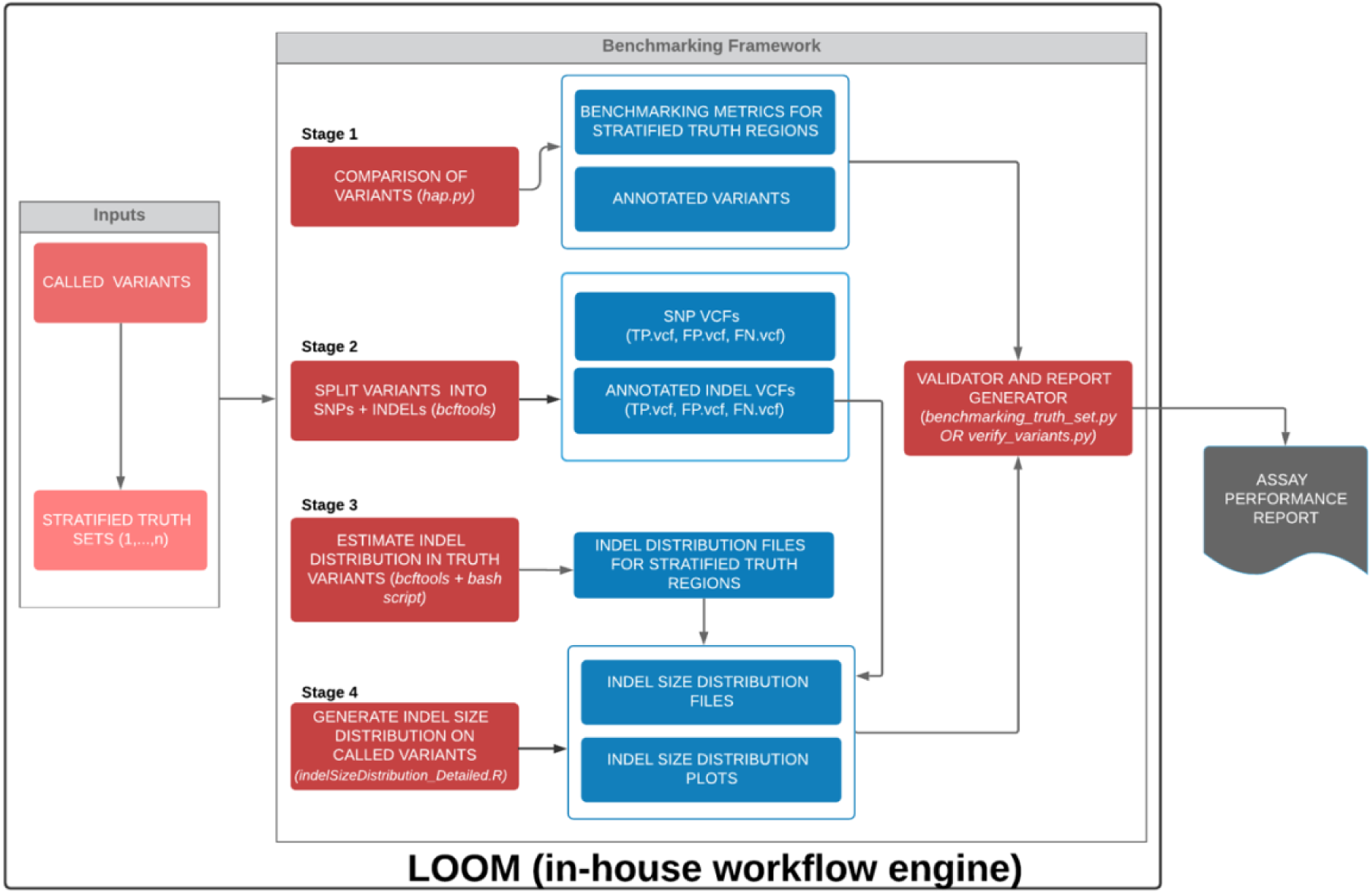
Schematic diagram of the benchmarking framework used in this study. All the stages in the benchmarking workflow have been dockerized. The docker images are available in DockerHub as specified in the Methods section.

The first step in the benchmarking process involves the comparison of input variants generated by the analytical pipeline with the benchmark variant calls within each region of interest. The variant calls are compared using hap.py (19, 20), which is capable of haplotype construction from individual genotype calls and is recommended by GIAB consortium and GA4GH. The variant comparison step is performed for each of the stratification or region of interest file specified as input, and hap.py generates a single output VCF file classifying the variant calls defined in the input and truth VCF files as either True Positive (TP), False Positive (FP) or False Negative (FN).

Step two in the benchmarking workflow splits the variant calls annotated using hap.py by variant type (SNPs and small INDELs) and by variant classification (TP/FP/FN). This step is executed within the workflow for each of the stratification or region of interest file specified. The VCF files are split by variant type using bcftools (21), and a bash script is used to further split the variant calls by the variant classification. This allows the workflow to generate the performance metrics for each of the variant types reported by the diagnostic assay.

Steps two and three of the benchmarking workflow (see Figure 1.) were used to generate a histogram of small INDELs by size. The bins used for INDEL size histograms were a. 1 base-pair, b. 2-5 base-pairs, c. 6-10 base-pairs, d. 11 – 20 base-pairs, e. 21 – 50 base-pairs, and f. Greater than 50 base-pairs. The R script - indelSizeDistribution_Detailed.R (code in

Additional File 1) then calculates the performance metrics of the assay for each of the INDEL size bins. The Python script – benchmarking_truth_set.py (Additional File 2) consolidates the benchmarking metrics previously obtained, calculates the NPA related metrics combining some of the bin size ranges (user preferred) for all reference samples provided.

In addition to benchmarking call sets for well-characterized reference samples published by the GIAB consortium, the benchmarking workflow allows for clinical laboratories to specify addition samples with clinically relevant variants as ground truths to estimate the analytical performance of the assay for specific variant types such as Δ*F*508 and IVS8-5T in CFTR panels. Python script – verify_variants.py (Additional File 3) accepts the ground-truth variant call sets to confirm the presence/absence of these variants in the VCF files generated by the variant calling pipeline. The details on the usage of the above scripts and associated README file are available in our public repository (also see Supplementary Materials).

Finally, the benchmarking workflow generates a comprehensive analytical validation report using all the provide benchmarking ground-truth call sets.

### Scalability and Reproducibility of Benchmarking workflow

The benchmarking workflow is designed to be repeatable and reproducible by using Docker containers for all software and bioinformatics components used within the workflow (see Table 6.). The workflow is distributed in human-readable data serialization format YAML v1.2, and the workflow can be readily executed using the workflow execution manager – LOOM (22). The workflow definition file – Benchmarking.yaml (see Supplementary Materials) can also be easily ported to Common Workflow Language (CWL) or Workflow Definition Language (WDL) formats and can be executed using workflow execution managers such as Toil (23, 24) and Cromwell (25).

**Table 6.**
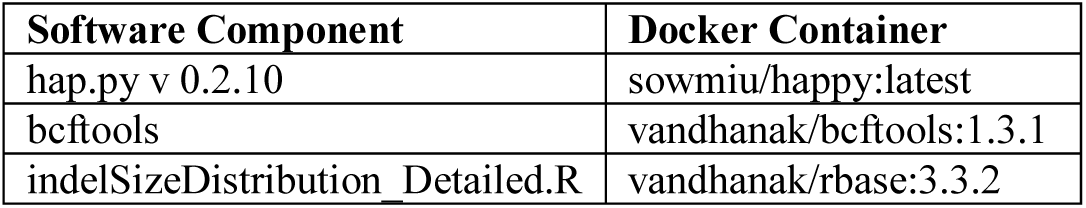
Docker containers and DockerHub repository location for each of the individual software components used in the benchmarking workflow.

| Software Component | Docker Container |
| --- | --- |
| hap.py v 0.2.10 | sowmiu/happy:latest |
| bcftools | vandhanak/bcftools:1.3.1 |
| indelSizeDistribution_Detailed.R | vandhanak/rbase:3.3.2 |

**Table 7.** GeT-RM sample ids and location of ground-truth variants in GRCh37 coordinates.

| GeT-RM Sample ID | Chromosome:Position | Truth Variant |
| --- | --- | --- |
| NA04408 | 15:91310152 | TATC -> T |
|  | 15:91310156 | T -> TA |
|  | 15:91310158 | A -> ATTC |
| NA14090 | 17:41276044 | ACT -> A |
| NA14170 | 13:32914437 | GT -> G |
| NA16658 | 10:43609103 | G -> T |

### Golden/ground-truth callsets

The golden/ground-truth sets for five reference and PGP genomes are currently available - NA12878 (CEPH family’s daughter), NA24143 (AJ mother), NA24149 (AJ father), NA24385 (AJ son), and NA24631(Chinese son) and these reference call sets were used in this benchmarking study. GIAB provides a high confidence regions file and a high confidence VCF file, and as recommended by GIAB, only the high confidence calls were used in the evaluation of the assay’s performance characteristics. The NIST versions and their corresponding FTP site locations used for the above samples in this study can be found in the Supplementary Material.

In addition to the GIAB reference samples, samples with known pathogenic germline variants (see Table 2.) for various inherited diseases/disorders were chosen from Genetic Testing Reference Materials Coordination Program (GeT-RM) (26-30)

### Stratification or Regions of Interest (ROI) BED files

Three stratification files were used to evaluate the performance characteristics of an inherited Whole Exome Sequencing (WES) assay.

1. Coding Exons for all known transcripts in RefSeq genes: RefSeq gene names, transcripts, and coordinates of all coding exons were obtained from the UCSC genome browser(31, 32).
2. Clinically relevant regions of the human genome: Clinically relevant regions were determined by intersecting coordinates of all known pathogenic variants reported in OMIM (33), ClinVar (34) and DECIPHER v9.28 (35) with the all exon regions (Coding and Non-Coding) file for RefSeq genes obtained from UCSC genome browser. The exonic coordinates were later extended by 20 base-pairs on either end to include canonical and non-canonical splice sites. Deep-intronic regions with pathogenic variants were added to the exonic regions to generate the final clinically relevant regions (BED) file.
3. Whole Exome regions file for RefSeq genes was obtained from UCSC genome browser. The exon regions were extended by 20 base-pairs on either end to include splice sites.

### Benchmarking metrics

Precision and recall are benchmarking metrics provided as output by hap.py. The true positives (TP), false positives (FP), and false negatives (FN) are counted as described by the developers of hap.py (20). Again, as explained by the authors of hap.py, precision and recall are calculated using the below formulae:

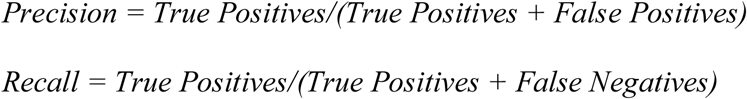

Other metrics reported by hap.py such as variants outside the high confidence truth set regions and transition or transversion SNP type can be found in the extended.csv files included in the Supplementary Materials.

The total number of bases per sample in a particular region of interest as specified by the corresponding bed file was computed using a bash command provided in the Supplementary Materials.

True negatives (TN) and Total Negatives are computed using the following:

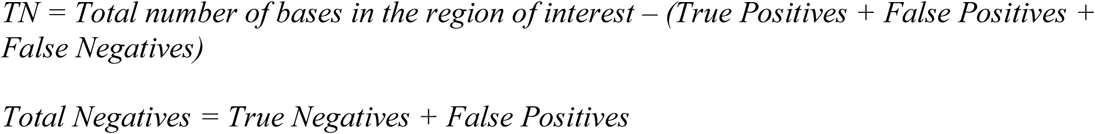

The Negative Percentage Agreement (NPA) or specificity as recommended by the FDA (36) is calculated using

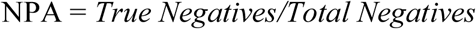

## List of abbreviations

NIST: National Institute of Standards and Technology
GIAB: Genome in a bottle consortium
SNPs: Single nucleotide polymorphisms
Indels: insertions/deletions
WES: Whole Exome Sequencing
NPA: Negative Percent Agreement
TN: True Negative
TP: True Positive
FN: False Negative
FP: False Positive
OMIM: public database containing the human genes, their genetic phenotypes and associations with genetic disorders (Online Mendelian Inheritance in Man)
DECIPHER: public database with genotypic and phenotypic data from ∼30,000 individuals
ClinVar: public database with information on the relationship between medically important variants and phenotypes.

## Declarations

Ethics approval and consent to participate

Not applicable

### Consent for publication

The authors declare that they have no competing interests.

### Availability of data and material

The datasets generated and/or analyzed during the current study are available in the GitHub repository - vandhanak/stanford-benchmarking-workflows.

### Authors’ contributions

VK designed and implemented the benchmarking workflow. SU and VK wrote the manuscript. ZN implemented the scripts to generate the performance assay report including the clinical variant validation. SU, SD, MP and EA conceived, designed and supervised the overall study. All authors read and approved the final manuscript.

## Acknowledgements

We thank Amin Zia for providing useful information during the initial phase of the benchmarking work. We thank Nathan Hammond and Issac Liao for the development of the in-house workflow engine “Loom” which was used to run the variant calling pipelines and the subsequent benchmarking workflows.

We are thankful to Chittaranjan Muthumalai for leading the automated benchmarking pipeline testing efforts and Jason Merker for useful discussions in terms of clinical relevance during the benchmarking process.

This study makes use of data generated by the DECIPHER community. A full list of centers who contributed to the generation of the data is available from http://decipher.sanger.ac.uk and via email from. Funding for the project was provided by the Wellcome Trust.

## Supplementary Materials

### Preparation of truth sets for exome regions

The NIST version and the ftp site used to download the original data for each of the GIAB samples (before preprocessing) used in this study are listed here.

NA12878

NIST v3.3:

ftp://ftp-trace.ncbi.nlm.nih.gov/giab/ftp/release/NA12878_HG001/NISTv3.3/NA12878_GIAB_highc onf_CG-IllFB-IllGATKHC-Ion-Solid-10X_CHROM1-X_v3.3_highconf.bed

ftp://ftp-trace.ncbi.nlm.nih.gov/giab/ftp/release/NA12878_HG001/NISTv3.3/NA12878_GIAB_highc onf_CG-IllFB-IllGATKHC-Ion-Solid-10X_CHROM1-X_v3.3_highconf.vcf.gz

NA24143

NIST v3.3:

ftp://ftp-trace.ncbi.nlm.nih.gov/giab/ftp/release/AshkenazimTrio/HG004_NA24143_mother/NISTv3.3/HG004_GIAB_highconf_CG-IllFB-IllGATKHC-Ion-10X_CHROM1-22_v3.3_highconf.bed

ftp://ftp-trace.ncbi.nlm.nih.gov/giab/ftp/release/AshkenazimTrio/HG004_NA24143_mother/NISTv3.3/HG004_GIAB_highconf_CG-IllFB-IllGATKHC-Ion-10X_CHROM1-22_v3.3_highconf.vcf.gz

NA24149

NIST v3.3:

ftp://ftp-trace.ncbi.nlm.nih.gov/giab/ftp/release/AshkenazimTrio/HG003_NA24149_father/NISTv3.3/HG003_GIAB_highconf_CG-IllFB-IllGATKHC-Ion-10X_CHROM1-22_v3.3_highconf.bed

ftp://ftp-trace.ncbi.nlm.nih.gov/giab/ftp/release/AshkenazimTrio/HG003_NA24149_father/NISTv3.3/HG003_GIAB_highconf_CG-IllFB-IllGATKHC-Ion-10X_CHROM1-22_v3.3_highconf.vcf.gz

NA24385

NIST v3.3:

ftp://ftp-trace.ncbi.nlm.nih.gov/giab/ftp/release/AshkenazimTrio/HG002_NA24385_son/NISTv3.3/HG002_GIAB_highconf_CG-IllFB-IllGATKHC-Ion-Solid-10X_CHROM1-22_v3.3_highconf.bed

ftp://ftp-trace.ncbi.nlm.nih.gov/giab/ftp/release/AshkenazimTrio/HG002_NA24385_son/NISTv3.3/HG002_GIAB_highconf_CG-IllFB-IllGATKHC-Ion-Solid-10X_CHROM1-22_v3.3_highconf.vcf.gz

NA24631

NIST v3.3.2:

ftp://ftp-trace.ncbi.nlm.nih.gov/giab/ftp/release/ChineseTrio/HG005_NA24631_son/NISTv3.3.2/GRCh37/HG005_GRCh37_highconf_CG-IllFB-IllGATKHC-Ion-SOLID_CHROM1-22_v.3.3.2_highconf_noMetaSV.bed

ftp://ftp-trace.ncbi.nlm.nih.gov/giab/ftp/release/ChineseTrio/HG005_NA24631_son/NISTv3.3.2/GRCh37/HG005_GRCh37_highconf_CG-IllFB-IllGATKHC-Ion-SOLID_CHROM1-22_v.3.3.2_highconf.vcf.gz

### Bash command to compute total number of bases in a region of interest

~~~
awk ’{a=$3-$2;print a}’ <Consolidated.bed> | paste -sd+ - | bc
~~~

In the above command, <Consolidated.bed> refers to GIAB original high confidence bed file for a sample intersected with the bed file of the region of interest such as coding exons, whole exome or clinically relevant gene regions. The user can use this command to calculate bases with their desired stratified region in the bed format which is required to compute metrics such as true negatives.

### Output files generated by Benchmarking workflow

Our benchmarking workflow generates the following output files:

~~~
1. <Output file common prefix>_<Sample ID>_CodingExons.vcf.gz
2. <Output file common prefix>_<Sample ID>_CodingExons.vcf.gz.tbi
3. <Output file common prefix>_<Sample ID>_CodingExons_counts.csv
4. <Output file common prefix>_<Sample ID>_CodingExons_counts.json
5. <Output file common prefix>_<Sample ID>_CodingExons_summary.csv
6. <Output file common prefix>_<Sample ID>_CodingExons_extended.csv
7. <Output file common prefix>_<Sample ID>_CodingExons_metrics.json
8. <Output file common prefix>_<Sample ID>_CodingExons_ConsoleOutput.txt
9. <Output file common prefix>_<Sample ID>_CodingExons_indelSizeDistribution.txt
10. <Output file common prefix>_<Sample ID>_CodingExons_indelSizeDistributionOnPlot.pdf
~~~

There is a final performance assay report generated in the form of a tab delimited file as below:

~~~
Final_benchmarking_metrics_*<current_date>.*txt
~~~

Another set of 10 files as seen above corresponding to the whole exome regions are generated.

The benchmarking framework generates the following intermediate files:

~~~
1. <Output file common prefix>_<Sample ID>_CodingExons_SNPs_TPonly.vcf.gz
2. <Output file common prefix>_<Sample ID>_CodingExons_SNPs_FPonly.vcf.gz
3. <Output file common prefix>_<Sample ID>_CodingExons_SNPs_FNonly.vcf.gz
4. <Output file common prefix>_<Sample ID>_CodingExons_INDELs_TPonly.vcf.gz
5. <Output file common prefix>_<Sample ID>_CodingExons_INDELs_FPonly.vcf.gz
6. <Output file common prefix>_<Sample ID>_CodingExons_INDELs_FNonly.vcf.gz
7. <Output file common prefix>_<SampleID>_CodingExons_indelDistribution.txt
~~~

Another set of seven files as seen above corresponding to the whole exome regions are generated.

## Supplemental Tables

**Table S1.** Benchmarking metrics for indels of different size ranges in NA12878 (truth set NIST v3.3, total bases = 13728555) for the regions within ∼7000 clinically relevant genes (as specified in Methods).

| Size of indels in NA12878 | Truth total | TP | FP | FN | TN | NPA | Precision | Recall |
| --- | --- | --- | --- | --- | --- | --- | --- | --- |
| 1-10 | 145 | 139 | 10 | 6 | 13728400 | 100 | 93.29 | 95.86 |
| 11-20 | 7 | 7 | 0 | 0 | 13728548 | 100 | 100 | 100 |
| 21-50 | 5 | 5 | 0 | 0 | 13728550 | 100 | 100 | 100 |
| All Indels | 156 | 150 | 10 | 6 | 13728389 | 100 | 93.75 | 96.15 |

**Table S2.** Benchmarking metrics for indels of different size ranges in NA24143 (truth set NIST v3.3, total bases = 12549224) for the regions within ∼7000 clinically relevant genes (as specified in Methods).

| Size of indels in NA24143 | Truth total | TP | FP | FN | TN | NPA | Precision | Recall |
| --- | --- | --- | --- | --- | --- | --- | --- | --- |
| 1-10 | 153 | 143 | 16 | 10 | 12549055 | 100 | 89.94 | 93.46 |
| 11-20 | 8 | 8 | 0 | 0 | 12549216 | 100 | 100 | 100 |
| 21-50 | 3 | 3 | 0 | 0 | 12549221 | 100 | 100 | 100 |
| All Indels | 163 | 153 | 16 | 10 | 12549045 | 100 | 90.53 | 93.87 |

**Table S3.** Benchmarking metrics for indels of different size ranges in NA24149 (truth set NIST v3.3, total bases = 12538042) for the regions within ∼7000 clinically relevant genes (as specified in Methods).

| Size of indels in NA24149 | Truth total | TP | FP | FN | TN | NPA | Precision | Recall |
| --- | --- | --- | --- | --- | --- | --- | --- | --- |
| 1-10 | 156 | 153 | 8 | 3 | 12537878 | 100 | 95.03 | 98.08 |
| 11-20 | 8 | 8 | 1 | 0 | 12538033 | 100 | 88.89 | 100 |
| 21-50 | 1 | 1 | 0 | 0 | 12538041 | 100 | 100 | 100 |
| All Indels | 163 | 161 | 9 | 3 | 12537869 | 100 | 94.71 | 98.16 |

**Table S4.** Benchmarking metrics for indels of different size ranges in NA24631 (truth set NIST v3.3, total bases = 12808688) for the regions within ∼7000 clinically relevant genes (as specified in Methods).

| Size of indels in NA24631 | Truth total | TP | FP | FN | TN | NPA | Precision | Recall |
| --- | --- | --- | --- | --- | --- | --- | --- | --- |
| 1-10 | 153 | 146 | 16 | 7 | 12808519 | 100 | 90.12 | 95.42 |
| 11-20 | 5 | 5 | 0 | 0 | 12808683 | 100 | 100 | 100 |
| 21-50 | 5 | 4 | 0 | 1 | 12808683 | 100 | 100 | 80 |
| All Indels | 162 | 154 | 16 | 8 | 12808510 | 100 | 90.59 | 95.06 |

**Table S5.** Benchmarking metrics on the number of indels of different size ranges in NA12878 (truth set NIST v3.3, total bases = 71152019) for the whole exome region s including non-coding exons, splice sites (+/-20 bp) and clinically relevant deep intronic regions.

| Size of indels in NA12878 | Truth total | TP | FP | FN | TN | NPA | Precision | Recall |
| --- | --- | --- | --- | --- | --- | --- | --- | --- |
| 1-10 | 5108 | 4704 | 781 | 404 | 71146130 | 100 | 85.76 | 92.09 |
| 11-20 | 209 | 194 | 13 | 15 | 71151797 | 100 | 93.72 | 92.82 |
| 21-50 | 52 | 47 | 5 | 5 | 71151962 | 100 | 90.38 | 90.38 |
| All Indels | 5318 | 4910 | 800 | 424 | 71145885 | 100 | 85.99 | 92.03 |

**Table S6.** Benchmarking metrics on the number of indels of different size ranges in NA24143 (truth set NIST v3.3, total bases = 65657646) for the whole exome regions including non-coding exons, splice sites (+/-20 bp) and clinically relevant deep intronic regions.

| Size of indels in NA24143 | Truth total | TP | FP | FN | TN | NPA | Precision | Recall |
| --- | --- | --- | --- | --- | --- | --- | --- | --- |
| 1-10 | 5168 | 4676 | 681 | 492 | 65651797 | 100 | 87.29 | 90.48 |
| 11-20 | 206 | 184 | 13 | 22 | 65657427 | 100 | 93.40 | 89.32 |
| 21-50 | 84 | 72 | 5 | 12 | 65657557 | 100 | 93.51 | 85.71 |
| All Indels | 5388 | 4878 | 700 | 526 | 65651542 | 100 | 87.45 | 90.24 |

**Table S7.** Benchmarking metrics on the number of indels of different size ranges in NA24149 (truth set NIST v3.3, total bases = 65597266) for the whole exome regions including non-coding exons, splice sites (+/-20 bp) and clinically relevant deep intronic regions.

| Size of indels in NA24149 | Truth total | TP | FP | FN | TN | NPA | Precision | Recall |
| --- | --- | --- | --- | --- | --- | --- | --- | --- |
| 1-10 | 5096 | 4578 | 628 | 518 | 65591542 | 100 | 87.94 | 89.84 |
| 11-20 | 188 | 167 | 17 | 21 | 65597061 | 100 | 90.76 | 88.83 |
| 21-50 | 68 | 62 | 5 | 6 | 65597193 | 100 | 92.54 | 91.18 |
| All Indels | 5290 | 4763 | 651 | 545 | 65591307 | 100 | 87.98 | 89.70 |

**Table S8.**
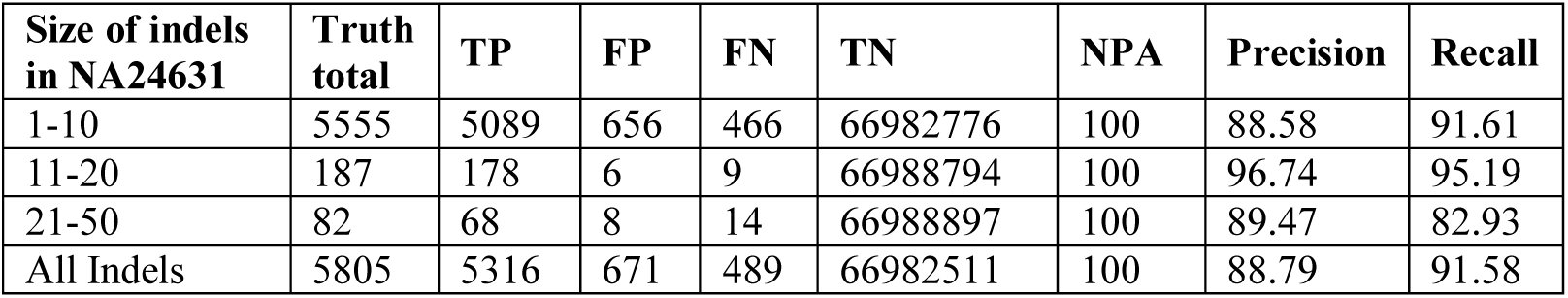
Benchmarking metrics on the number of indels of different size ranges in NA24631 (truth set NIST v3.3, total bases = 65657646) for the whole exome regions including non-coding exons, splice sites (+/-20 bp) and clinically relevant deep intronic regions.

| Size of indels in NA24631 | Truth total | TP | FP | FN | TN | NPA | Precision | Recall |
| --- | --- | --- | --- | --- | --- | --- | --- | --- |
| 1-10 | 5555 | 5089 | 656 | 466 | 66982776 | 100 | 88.58 | 91.61 |
| 11-20 | 187 | 178 | 6 | 9 | 66988794 | 100 | 96.74 | 95.19 |
| 21-50 | 82 | 68 | 8 | 14 | 66988897 | 100 | 89.47 | 82.93 |
| All Indels | 5805 | 5316 | 671 | 489 | 66982511 | 100 | 88.79 | 91.58 |

## Supplemental Data files for

### Benchmarking workflows to assess performance and suitability of germline variant calling pipelines in clinical diagnostic assays

The benchmarking workflow file and relevant scripts (listed below as additional files) and all output files for five GIAB samples per stage are available in our public repository: vandhanak/stanford-benchmarking-workflows

### Supplemental Files

1. indelSizeDistribution_Detailed.R
2. benchmarking_truth_set.py
3. verify_variants.py

